# Direct observation of the evolution of cell-type specific microRNA expression signatures supports the hematopoietic origin model of endothelial cells

**DOI:** 10.1101/2022.11.23.517662

**Authors:** Ana E. Jenike, Katharine M. Jenike, Kevin J. Peterson, Bastian Fromm, Marc K. Halushka

## Abstract

The evolution of specialized cell-types is a long-standing interest of biologists, but given the deep time-scales very difficult to reconstruct or observe. microRNAs have been linked to the evolution of cellular complexity and may inform on specialization. The endothelium is a vertebrate specific specialization of the circulatory system that enabled a critical new level of vasoregulation. The evolutionary origin of these endothelial cells is unclear. We hypothesized that Mir-126, an endothelial cell-specific microRNA may be informative.

We here reconstruct the evolutionary history of Mir-126. Mir-126 likely appeared in the last common ancestor of vertebrates and tunicates, a species without an endothelium, within an intron of the evolutionary much older EGF Like Domain Multiple (Egfl) locus. Mir-126 has a complex evolutionary history due to duplications and losses of both the host gene and the microRNA. Taking advantage of the strong evolutionary conservation of the microRNA among Olfactores, and using RNA in situ hybridization (RISH), we localized Mir-126 in the tunicate *Ciona robusta*. We found exclusive expression of the mature Mir-126 in granular amebocytes, supporting a long-proposed scenario that endothelial cells arose from hemoblasts, a type of proto-endothelial amoebocyte found throughout invertebrates.

This observed change of expression of Mir-126 from proto-endothelial amoebocytes in the tunicate to endothelial cells in vertebrates is the first direct observation of the evolution of a cell-type in relation to microRNA expression indicating that microRNAs can be a prerequisite of cell-type evolution.

**Research Highlights:**

- direct observation of cell-type evolution
- high conservation of sequence enables for simple RISH experiment of expression
- Mir-126 follows the evolution of hematopoetic cells to endothelial cells

## 1 INTRODUCTION

Increasing cellular diversity is a hallmark of metazoan evolution (Deline et al., 2018). Naturally, therefore it has been a focus of biologists from all disciplines to understand just how many cell types there are, how to define them and, most importantly, how new cell types have evolved through evolution (see (Arendt, 2008; Arendt et al., 2016; Kin et al., 2015; Wang et al., 2021). All views and models on these sometimes controversially discussed subjects, however, agree that during animal evolution a substantial increase of morphological complexity facilitated by an expansion of distinct cell-types can be observed at several key points during animal evolution, in particular at the origin of our very own clade, the Vertebrata (Deline et al., 2018; Musser et al., 2021; Valentine et al., 1994; Wagner, 2014). Several innovative technologies have arisen that allow for the dissection of the molecular basis of cell types and the composition of corresponding gene regulatory networks including bulk RNA-seq on cultured cell-lines (Kin et al., 2015) as well as single cell sequencing focused studies that have allowed extensive insights on cell types and cell-population differences with clear applications to diseases (Jagadeesh et al., 2022). It is clear that novel cell types do not originate de novo, but like in a genealogical tree, descend from related and evolutionarily older cell-types.

Nonetheless, when comparing animals for their cell-type numbers, it is very hard to find genomic features, or elements that would correlate with the differences in cellular complexity. These observations are widely known as the C-value paradox (for genome size) (Cavalier-Smith, 1978) and the G-value paradox (number of protein coding games) (Hahn & Wray, 2002), respectively. This lack of correlation indicates a combinatorial, or regulatory origin of cellular complexity and it has been proposed to originate in the non-coding part of genomes (Taft et al., 2007). Indeed, the advent of novel microRNAs, ∼22 nucleotide short non-coding post-transcriptional gene regulators (Bartel, 2018), strongly correlates both with increasing complexity as found in several metazoan clades including vertebrates and cephalopod mollusks (Deline et al., 2018; Grimson et al., 2008; Heimberg et al., 2008; Sempere et al., 2006; Tanzer & Stadler, 2006; Wheeler et al., 2009; Zolotarov et al., 2022). Their loss also accompanies morphological simplification as seen in, for example, parasitic species (Fromm et al., 2013).

The discovery of microRNAs pre-dated Next Generation Sequencing or single-cell methods (Lee et al., 1993), and hence expression of microRNAs was initially characterized at the organ or tissue level (Lagos-Quintana et al., 2002; Landgraf et al., 2007; Volinia et al., 2006). These studies revealed that microRNA evolution and the establishment of tissue identities were closely coupled in bilaterian evolution (Christodoulou et al., 2010). From cell-culture based bulk data, the cellular expression levels for many microRNAs were established (Haider et al., 2014; Landgraf et al., 2007; McCall et al., 2011). More recently, extensive cellular microRNAome data became available that established a quantitative framework for cellular microRNA expression (de Rie et al., 2017; Lorenzi et al., 2021; McCall et al., 2017; Patil et al., 2022). microRNAs essentially fall in two classes: ubiquitously expressed microRNAs, i.e. microRNAs found in many cell-types and tissues such as LET-7 or MIR-21, and cell-type specific microRNAs such as MIR-1, MIR-122, MIR-124, and MIR-451, which are exclusively found in only one or few cell types (Patil et al., 2022). This latter group indicates a specific function of these microRNAs that may impact on the speciation of cell types. A detailed and comparable annotation record for individual microRNAs across species can untangle the evolutionary origin of these cells types and the roles microRNA played. MicroRNAs can be either a prerequisite for the evolution of the particular cell type, or a consequence that helped to canalize the gene expression patterns of the protein coding genes (Kosik, 2010). Given that most microRNAs do not initiate specification, but instead help lock-in a differentiation pathway that is unique for a given cell type (Makeyev & Maniatis, 2008), most authors argue that microRNAs help canalize the developmental process by conferring robustness to the specification process (Ebert & Sharp, 2012; Hornstein & Shomron, 2006; Peterson et al., 2009; Posadas & Carthew, 2014).

One of the major evolutionary events in animal evolution, and of substantial microRNA expansion, is the origin of vertebrates around 500 million years ago. Numerous new cell types and tissues characterize vertebrates (Heimberg et al., 2008; Trainor, 2013; York & McCauley, 2020) including the endothelium, a key adaptation of the circulatory system that enabled a critical new level of vasoregulation. Endothelial cells are a specialized cell type that line blood and lymphatic vessels in all vertebrate species (Monahan-Earley et al., 2013). These cells play an active role in many biological processes in the body, including trafficking of blood and nutrients, blood fluidity, providing a vascular barrier, and immunity (Khazaei et al., 2008; Lerman & Zeiher, 2005; Vita, 2011). Interestingly, while it is clear that endothelial cells first evolved in vertebrates, their evolutionary origin and ancestral cell-type is unclear (Monahan-Earley et al., 2013). One hypothesis is that endothelial cells arose from mesenchymal cells that can differentiate into endothelial cells and smooth muscle cells (Yamashita et al., 2000). Alternatively, endothelial cells possibly evolved from multipotent blood precursor cells (known as hemangioblasts) (Sabin, 1920) that are related to ancient blood-derived cells called amebocytes (Monahan-Earley et al., 2013; Munoz-Chapuli et al., 2005). These cells are also thought to be the precursors of thrombocytes indicating that endothelial cells and thrombocytes could be sister cell types (Levin, 2019). Further, in annelids, mollusks, and echinoderms, amebocytes can line the walls of blood vessels, suggesting that this cell type could be a form of proto-endothelium (Harrison & Foelix, 1999). Some main proteins investigated to understand endothelial evolution are vascular endothelial growth factor (VEGF), its receptor (VEGFR), von Willebrand factor (vWF) and vascular cell adhesion molecule 1 (VCAM1) (Holmes & Zachary, 2005; Lenting et al., 2010; Munoz-Chapuli et al., 2005). However, the evolution of these proteins either long predates the evolution of endothelial cells (VEGF, VEGFR, vWF), or exactly co-occurs (VCAM1) with vertebrates, leading to a gap in our knowledge of vascular cellular evolution.

We here take advantage of the presence of an endothelial cell enriched microRNA (Mir-126) in *Ciona robusta*, a member of the Tunicata and as such the sister taxon to the vertebrates. In vertebrates including teleost fish and mammals (Figure 1A-D), Mir-126 is known to be nearly exclusively and strongly expressed in endothelial cells (Fish et al., 2008; Wienholds et al., 2005), but also lowly expressed in thrombocytes (megakaryocytes and platelet particles) and possibly at even lower levels in other hematologic cells (Figure 1E) (McCall et al., 2011; Patil et al., 2022). Mir-126 is a key regulator of endothelial-specific transcripts through both direct (e.g., VCAM1) as well as indirect (e.g., VEGF) regulation impacting the process of vasculogenesis (Harris et al., 2008; Nicoli et al., 2010; Wang et al., 2008). Knockout of Mir-126, but not of the hosting protein-coding gene *Egfl7*, results in low rates of angiogenesis and no vascular healing (Kuhnert et al., 2008; Wang et al., 2008). Although tunicates lack a true endothelium, Mir-126 expression in *C. robusta* increases from larval to adult stages (Shi et al., 2009) suggesting a role in the specification of adult-specific cell types, but its localization is unknown. Nonetheless, given the expression profile of Mir-126 in vertebrates, we hypothesized that the expression of Mir-126 in tunicates might shed light on the origin of endothelial and thrombocyte cell types. Using RNA-in-situ-hybridization (RISH), we find that Mir-126 in the tunicate *Ciona robusta* is exclusively expressed in granular amebocytes, an invertebrate hemoblast cell, supporting the long-proposed scenario that endothelial cells arose from hemoblasts and are related to megakaryocytes, thrombocytes, and platelets.

**Figure 1.**
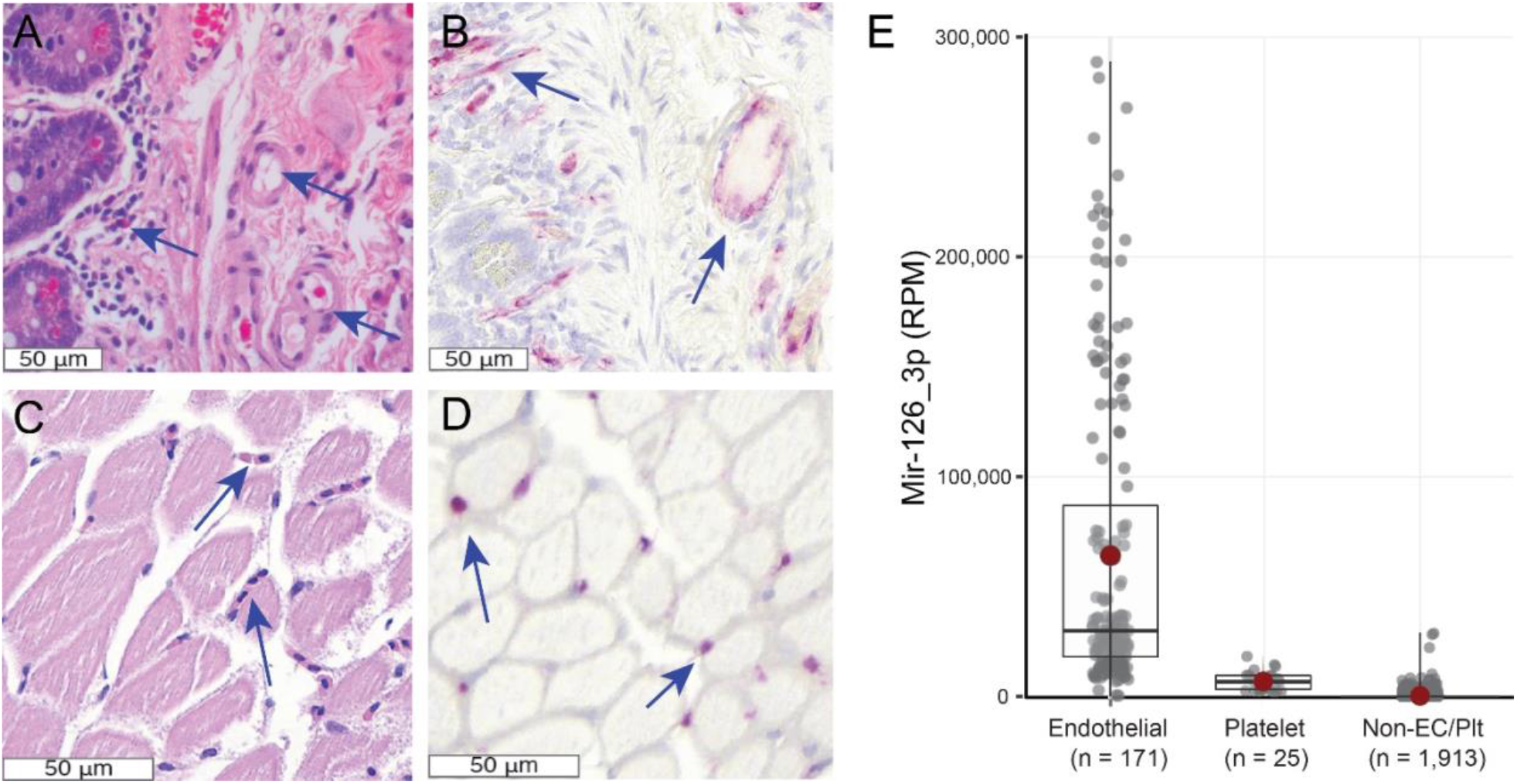
Mir-126_3p localization and expression. **A**. Human intestinal tissue. The blue arrows highlight vascular spaces. **B**. A Mir-126_3p RISH probe highlights endothelial cells along blood vessels. **C**. Skeletal muscle of a zebrafish (*Danio rerio*) with capillaries identified by blue arrows. D. The same Mir-126_3p probe highlights capillary endothelial cells. **E**. Mir-126_3p is highly expressed among human endothelial cells, modestly expressed among platelets and is essentially absent across 182 cell types (Non-EC/Plt). Red dot is average, bar is median. Data from (Patil et al., 2022)

## 2 MATERIAL AND METHODS

### 2.1 Nomenclature of *Ciona robusta*

The nomenclature concerning *Ciona robusta* is complicated. It was officially recognized as a separate species to *Ciona intestinalis* in 2015, based on work by Hoshino and Tokioka (Hoshino & Tokioka, 1967), followed by Brunetti et al. (Brunetti et al., 2015). Thus, much of the historical *C. intestinalis* literature may be referring to the *C. robusta* tunicate. In this manuscript, *C. intestinalis, C. robusta*, and tunicate terms are used as accurately as possible.

### 2.2 MIR-126 sequence acquisition and synteny analyses

Precursor sequences of MIR-126 representatives of the tunicates (*Ciona intestinalis / robusta*), cyclostomes (the sea lamprey, *Petromyzon marinus* and the inshore hagfish, *Eptatretus burgeri*), sharks (the ghostshark, *Callorhinchus milii*) and teleost species representing Arctinopterygii (ray-finned fish: spotted gar, *Lepisosteus oculatus* and zebrafish, *Danio rerio*) and Sarcopterygii including the coelacanth *Latimeria chalumnae*, the chicken, *Gallus gallus*, house mouse, *Mus musculus* and human, were downloaded from MirGeneDB.org (Fromm et al., 2022). If processing variants were annotated, only variant 1 was included in alignments. In addition, MIR-126 was blasted against available tunicate genomes and genomic traces on NCBI. Synteny analyses and orthologue / paralogue assignment were conducted as described before (Fromm et al., 2022). Briefly, using Ensembl or UCSC, respectively, all annotated microRNA loci were checked whether they resided in a protein coding gene with similarities to *EGLF* and whether Notch or *RGL* orthologues could be identified up or downstream of the microRNA.

### 2.3 Mir-126 read counts

Expression of Mir-126, based on reads per million (RPM) values across 171 endothelial cells, 25 platelets and 1,913 cells that were not endothelial cells or platelets were obtained from the ABC of cellular microRNAome Track Hub at the UCSC Genome Browser (Patil et al., 2022).

### 2.4 *Ciona robusta* acquisition and dissection

The *C. robusta* were collected fresh from the Southern California coast (M-REP) and overnight shipped to Johns Hopkins University where they were kept in a 20 gallon tall saltwater tank at 33.9 ppt. A sacrificing reagent was made with 2% isopropanol (VWR, cat no. 0918-1L) and TBS at 2% (Sigma Aldrich, MFCD00132476). The tunicates were sacrificed in isopropanol and refrigerated at 4°C for 15 minutes. The tunicates were then fixed in formalin (10% neutral buffered) (Figure S1).

After 24 hours, four tunicates were removed from fixation. Two tunicates were sagittally cut and two tunicates were vertically cut. Sections were placed in tissue cassettes, and further fixed for 68 hours due to high cellulose and high-water content. After 48 hours, two more tunicates were taken out of formalin and dissected sagittally. These sections were placed in tissue cassettes, and fixed for an additional 68 hours. For two tunicates, the tunics were dissected away, and the remainder of the tunicate was fixed for 24 hours. The tissue was then sectioned, placed in a tissue cassette, and further fixed for 48 hours.

Two small tunicates, under 3cm, were taken whole into tissue cassettes. The larger of the two tunicates had its tunic pierced 7 times along the vertical plane with a wet blade, to allow easier formalin penetration. The tunicate was then placed in a tissue cassette and fixed for 48 hours before processing. The smaller of the two tunicates was taken whole into a tissue cassette and fixed for 48 hours before processing.

Each tissue was paraffin embedded and hematoxylin and eosin (H&E) stained slides were generated on each block. A subset of slides were stained for Periodic Acid Schiff (PAS) to highlight glycogen.

### 2.5 Siphon regrowth experiment

To test the expression of Mir-126 during regrowth, siphons were amputated from six tunicates (2-3 mm above the apex of the siphons), after which the tunicates were returned to the salt water tank, in separated baskets. At time points 48 hours and 96 hours post-amputation, tunicates (N = 3 at 48 hours and 2 at 96 hours) were sacrificed in isopropanol at 4°C for 15 minutes. One tunicate did not survive to 96 hours. Sections were taken of the tissue, horizontal to the plane of amputation, and fixed for 48 hours in tissue cassettes.

### 2.6 RNA in situ hybridization staining for Mir-126_3p

After evaluating H&E slides, 6 blocks were selected for their diversity of organs shown. Staining for Mir-126 on human tissue, *D. rerio* tissue and *C. robusta* was performed using the same ACD Biosciences manual red microRNA *in situ* hybridization kit (ACD, Cat no. 322300, 322360, 310091) across all species. Two positive controls were used, human small intestine and a cross section of *D. rerio*. Staining was performed per the ACD recommended protocol. The tunicate staining followed the ACD protocol with the exception of protease antigen retrieval and AMP5 incubations, which were shortened to 7.5 minutes.

## 3 Results

### 3.1 The evolution of Mir-126 and the host EGFL7/8 locus

In human and most vertebrates, there is a single Mir-126 gene lying intragenically in its host gene *Egfl7* (Figure 2). However, jawed vertebrates are characterized by the possession of two *Egfl* paralogues, *Egfl7* and *Egfl8*, which are descended from a single ancestral *Egfl* gene present throughout most metazoans. Vertebrates underwent a single whole genome duplication (WGD) before the last common ancestor (LCA) of jawless (cyclostomes) and jawed (gnathostomes) fish (known as 1R). After this vertebrate LCA, but before the LCA of gnathostomes, the jawed fish underwent a second WGD (known as 2R). *Egfl7* and *Egfl8* are the products of this second WGD as they are present on the sub-genomes generated from 2R (Lamb, 2021; Simakov et al., 2020) (Figure 1). In the bony fish lineage *Egfl8* lost its embedded *Mir-126* locus, and a single Mir-126 locus was retained within *Egfl7*. However, in the Chondrichthyes, both Mir-126 loci are retained, one (Mir-126-P4) within *Egfl8*, and a second locus orthologous to the human Mir-126 gene (Mir-126-P2) that now lies intergenically as the host *Egfl7* gene was lost (Figure 2). Thus, similar to what is seen with other ancient microRNAs (e.g., Mir-193, (Campo-Paysaa et al., 2011)), either the host gene and/or the microRNA can have separate evolutionary trajectories in terms of gene retention versus gene loss especially following WGDs. Here, *EGFL7* and Mir-126-P2 are retained in human, whereas *Egfl8* and Mir-126-P4 are retained in shark. Furthermore, although *EGFL8* is retained in human, it lost its embedded Mir-126-P4 gene. On the other hand, although Mir-126-P2 is retained in shark, it is orphaned due to the loss of the *Egfl7* gene.

**Figure 2.**
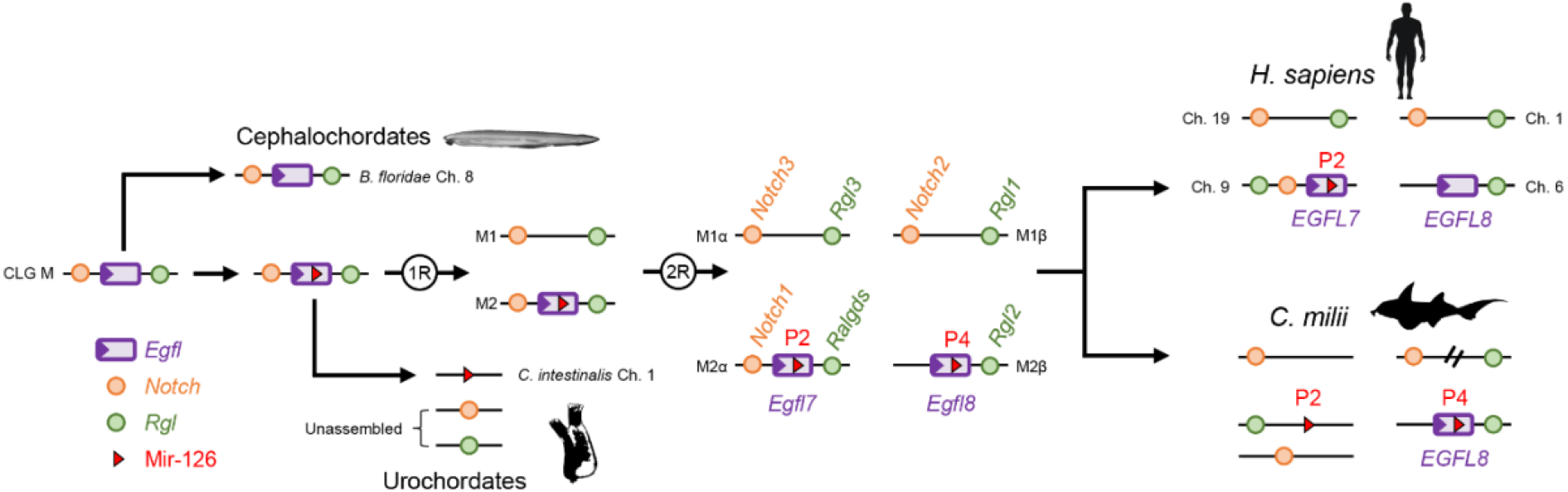
The evolutionary history of Mir-126 and its host protein-encoding gene *Egfl7/8*. The phylum Chordata includes three primary subgroups, the cephalochordates (or amphioxus, including the species *Branchiostoma floridae*), the tunicates that includes the ascidians or sea squirts (e.g., *Ciona intestinalis*) and the vertebrates, represented by human (*H. sapiens*) and the elephant shark (*C. milii*). Comparative genomic studies indicate that the chordate last common ancestor (LCA) had ∼17 chromosomes or chordate linkage groups (CLG) (Lamb, 2021; Simakov et al., 2020). The ancestral single copy *Egfl* gene (purple) resided on CLG M and was syntenic with two other genes, *Notch* (orange) and *Rgl* (green). These three genes remain syntenic in *B. floridae* and are found on chromosome 8, the descendant chromosome of CLG M in amphioxus (Simakov et al., 2020). After the split from cephalochordates, but before the split between the tunicates and vertebrates, the miRNA Mir-126 evolved within an intron of the ancestral *Egfl* gene (red arrowhead). Within the tunicate lineage the host *Egfl* gene was lost, orphaning the embedded Mir-126, which is now found on chromosome 1 in the ascidian *C. intestinalis*. After this LCA, jawed vertebrates underwent two whole genome duplication events (1R and 2R), ultimately generating four separate sub-genomes (1α, 1β, 2α and 2β). After 1R, sub-genome 1 lost both the *Egfl* paralogue as well as the embedded miRNA-126, but both genes are retained on sub-genome 2. Following 2R, the remaining single-copy *Egfl* gene, as well as the miRNA, duplicated into two separate paralogues: M2α hosts *Egfl7* and Mir-126-P2 whereas M2β hosts *Egfl8* and Mir-126-P4. In the evolution of bony fish after the split from the Chondrichthyes, *Egfl8* lost the Mir-126-P4 gene, and most taxa outside of mammals and sharks also lost the *Egfl8* gene. On the other hand, the elephant shark lost the *Egfl7* gene, but retains both ancestral copies of Mir-126 on separate genomic contigs.

Beyond the loss of *Egfl7* loss in Chondrichthyes, the loss of *Egfl* loci is quite common in chordate evolution. For example, teleosts lost both the ancestral *Egfl8* locus as well as one of the two *Egfl7* loci generated during the teleost-specific WGD (3R), but yet retained both copies of Mir-126-P2 generated from this 3R event. Tunicates, the nearest invertebrate relative of the vertebrates, are particularly interesting in this regard. Although no known tunicate retains the ancestral *Egfl7/8* host gene (Figure 2), representative species of two of the three major sub-taxa (the “ascidians” or sea squirts and the thaliaceans or salps) possess a single Mir-126 locus (Hendrix et al., 2010; Norden-Krichmar et al., 2007; Wang et al., 2017) (Figure 3); only appendicularians with their dramatically reduced genome (Bliznina et al., 2021) – including the loss of numerous conserved microRNAs (Fu et al., 2008) – lack the Mir-126 locus.

**Figure 3.**
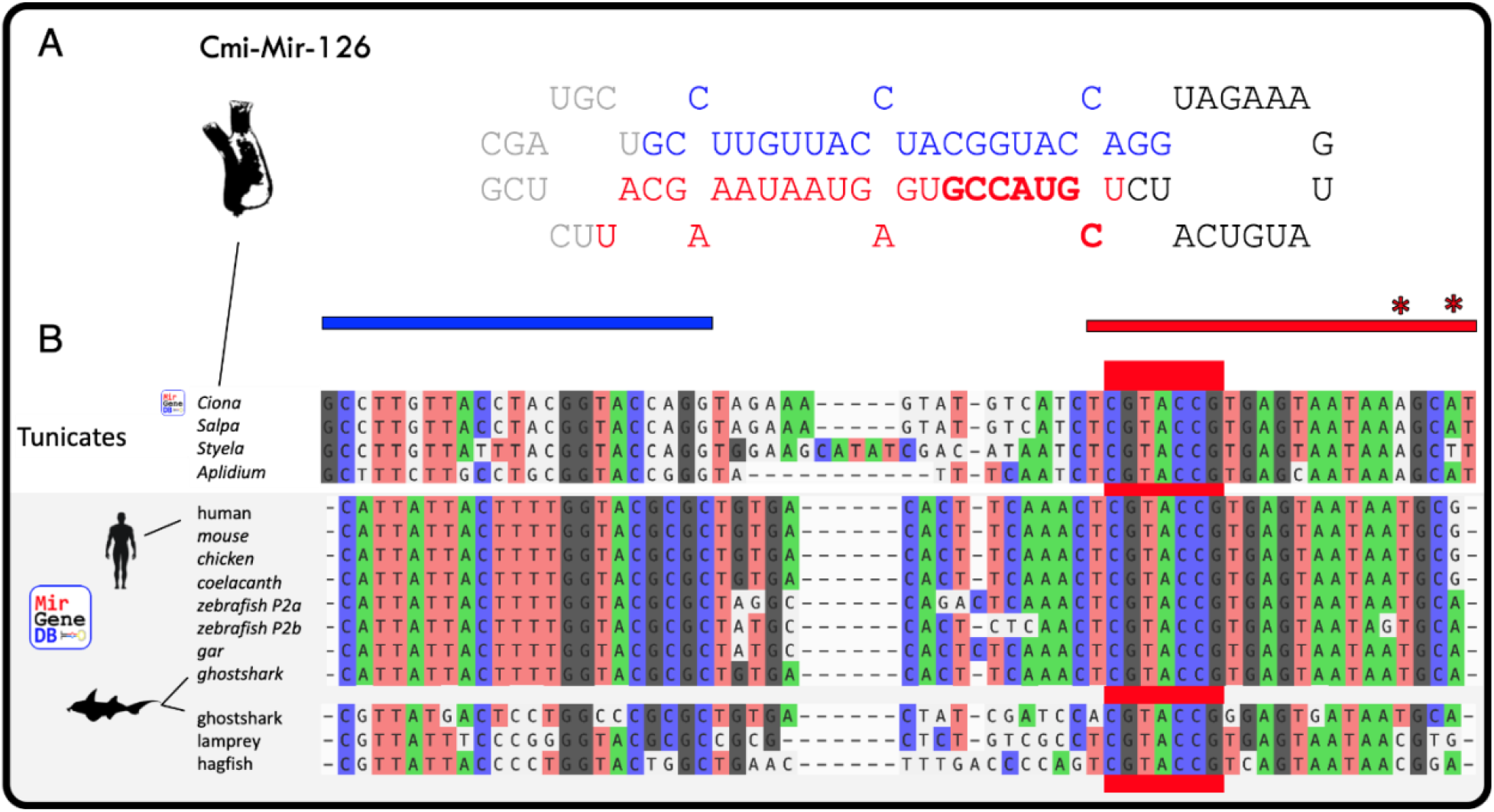
Genomic conservation of the Mir-126 locus in representative species from MirGeneDB and herein identified orthologues in tunicates. **A**. The hairpin loop structure and sequence in *C. intestinalis*. The 5p passenger strand is highlighted in blue and the 3p mature strand is highlighted in red. **B**. Sequence conservation across species ranging from early vertebrates such as sharks to eutherian mammals. There is strong evolutionary conservation of Mir-126_3p across 500 million years of evolution, with no changes to the CGUACCG seed sequence across any species and only two nucleotide changes between *C. intestinalis* and *H. sapiens* (position 18 and 22, asterisks). The 5p strand is well conserved among vertebrate species, but differs by 7 nucleotides between *C. intestinalis* and *H. sapiens*.

### 3.2 Mir-126 sequence is highly conserved in Olfactores

An exhaustive analysis of microRNA evolution across Metazoa (Fromm et al., 2020) reveals that the MIR-126 family is restricted to the Olfactores, the group that includes the tunicates and the vertebrates (Delsuc et al., 2008) that diverged from one another nearly 550 million years ago (Delsuc et al., 2018). As expected, the Mir-126 locus is highly conserved with the tunicate representatives showing high sequence similarity across the whole precursor and, as expected, higher sequence divergence in the star (5p) arm relative to the mature (3p) arm. Indeed, there are only two nucleotide difference in the mature Mir-126 sequence between *C. intestinalis* and *H. sapiens* (Figure 3). Within vertebrates, higher rates of sequence evolution are observed in the paralogue group found only in the cyclostomes and shark relative to the P2 group, as expected from comparisons of all ancestral microRNA genes in vertebrates (Peterson et al., 2022). Altogether, these patterns are typical conservation patterns of microRNAs (Fromm et al., 2015) and raised the interesting possibility that existing RISH probes for human Mir-126_3p could work for all species, including the tunicate.

### 3.3 Localization of Mir-126 in C. robusta to granular amebocytes

The strong conservation of the mature arm of the microRNA led us to test the suitability of RISH probes from human in the tunicate. For this, we successfully validated the RISH method in human and *D. rerio* tissues indicating specific endothelial cell staining (Figure 1). We then stained *C. robusta* tissues and identified Mir-126_3p localization. Across all organs and tissues, the blood-based granular amebocyte was the only cell type to stain positive for Mir-126_3p (Figure 4A, Figure S2). Blood cells in *C. robusta* cluster by type in both the lymph node/hematopoietic organ and in circulation. This allowed for identification of Mir-126 positive cells through comparing the cells to a serially adjacent H&E stained slide (Figure 4B). In some cases, granular amebocytes can be seen adhering to the internal side of blood lacunae, a known functional activity of these cells. Mir-126 positive granular amebocytes were seen frequently in the lymph node/hematopoietic organ (∼20% of all cells), the blood lacunae, and rarely within the test (tunicate) matrix. Importantly, all epithelial cells, muscle cells, sex cells, and specialty cells of the tunicate were negative for Mir-126_3p staining, indicating specific Mir-126 expression in amebocytes. Granular amebocytes are ∼8% of the total blood cell population and are noted histologically and by ultrastructural examination as having electron-dense vanadium deposits and a zone of microtubules (Rowley, 1982). This blood cell type is capable of penetrating the blood lacunae walls in response to injury. It can express complement C3, a natural killer cell gene homolog (CiCD94-i), and has phagocytic capabilities (Pinto et al., 2003; Zucchetti et al., 2008). It is similar in many respects to the amebocytes of other invertebrates that coat the walls of blood lacunae.

**Figure 4.**
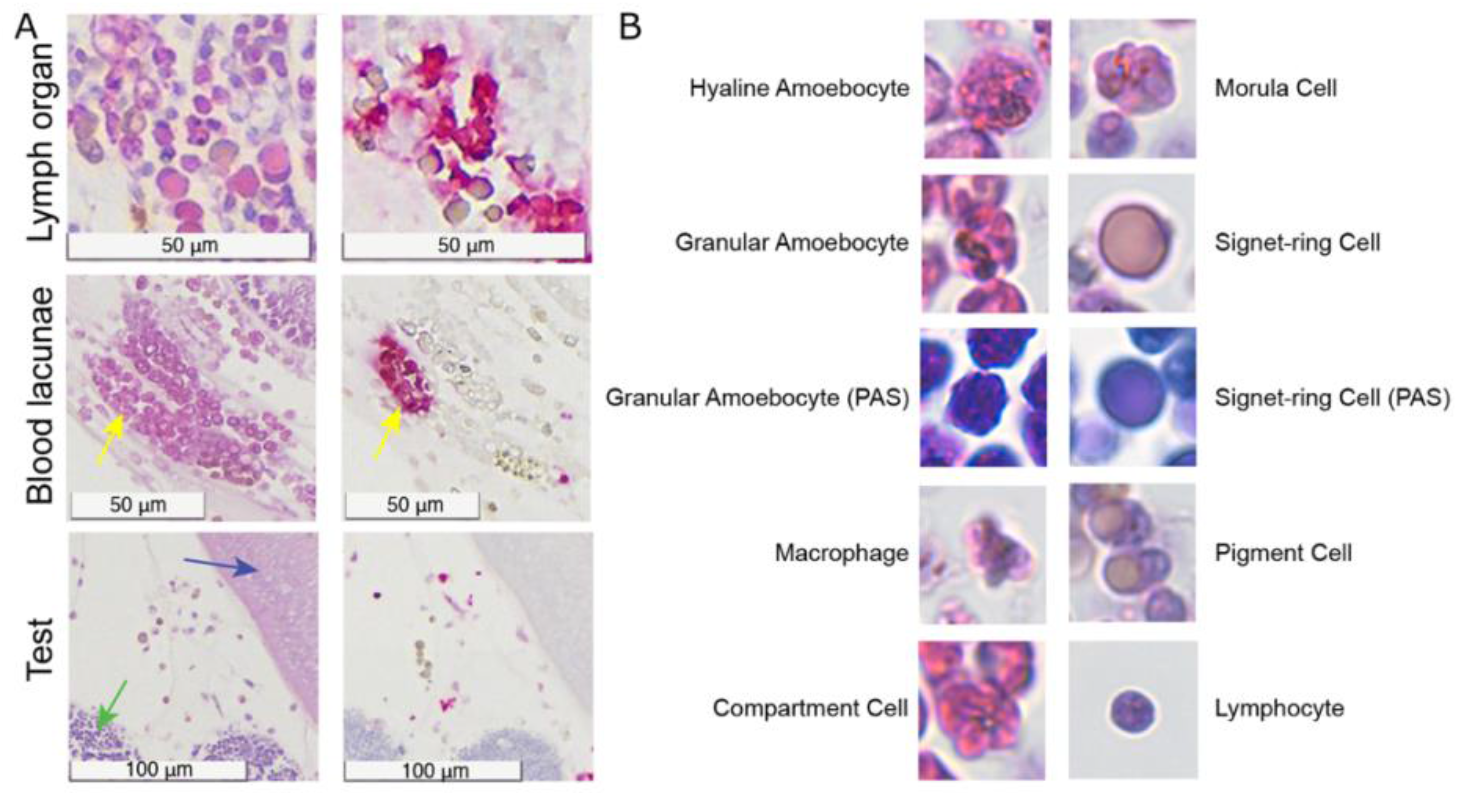
Mir-126_3p localization. **A**. Mir-126_3p is expressed exclusively in granular amebocytes. This is noted in collections of mixed blood cells in the lymph organ, in the blood lacunae (highlighted by the yellow arrows), or in the test (ground substance of the cuticle). The blue arrow points to intestinal epithelium and the green arrow to the pyloric gland, which did not express Mir-126. **B**. Histologic representations of each blood cell type based on (Millar, 1948). False staining and blush were noted in the cuticle, ovary, and neural gland (Figure S2).

### 3.4 Correlation of Mir-126 expression and granular amebocyte infiltrate in a siphon injury model

Ciona are known to induce a granulation tissue-type response to injury (Nicolo et al., 2016) which multiple blood cells including granular amebocytes, invade into the test. We reasoned that a siphon injury model could be useful in confirming the discovery of Mir-126 in granular amebocytes and determine if any other cell types activated expression of Mir-126 in this perturbed state. Six *C. robusta* underwent siphon amputation, with harvesting at 2 (n=3) and 4 (n=2) days post injury. We noted a ∼5-fold increase in blood cells present in the siphon test at 48 hours post injury (139±18 cells per 5,625 μm^2^) relative to a healthy comparable area (26±12 cells) at day 0. By 96 hours post injury, the cellular infiltrate decreased to 32±11 cells. Granular amebocytes, were a lesser constituent of the infiltrate being only 20±4 cells and 6±2 cells at 48 and 96 hours respectively. Mir-126+ cell counts at these same time points were 11±6 and 3±2 cells capturing roughly half of the granular amebocyte infiltrate and suggesting a general upregulation of Mir-126 in other infiltrating cell types was unlikely (Figure 5). By histology, signet ring cells, a cell type not noted to express Mir-126, made up a dominant percentage of the infiltrating cells.

## 4 DISCUSSION

The evolution of cellular complexity is a recurrent interest of evolutionary biologists and cell biologist alike. However, the study of cellular evolution faces several methodological challenges. Among others, a main challenge of the field is to agree on the total number of distinct cell types and, hence, to establish a reproducible approach to distinguish cell types morphologically, transcriptionally or by other means. Recently, the integration of both biochemical as well as transcriptional information has allowed for much better understanding of neuronal cell types (e.g., (Zeisel et al., 2018)) allowing for a more complete picture of, for example, brain evolution through deep time (Hain et al., 2022; Woych et al., 2022). However, despite these remarkable technological developments and biological insights, they often suffer from the *a priori* exclusion of the non-coding genome, in particular the expression of microRNAs. Further, when microRNAs are considered as markers for cell-type identity, a difficulty immediately arises in that the evolution of the microRNA (whether, for example, MIR-1 or MIR-122) is coincident with the advent of the cell type (the striated muscle of bilaterians or the hepatocyte of vertebrates, respectively) as both the microRNA and the cell type co-evolve at the same point in metazoan evolution.

**Figure 5.**
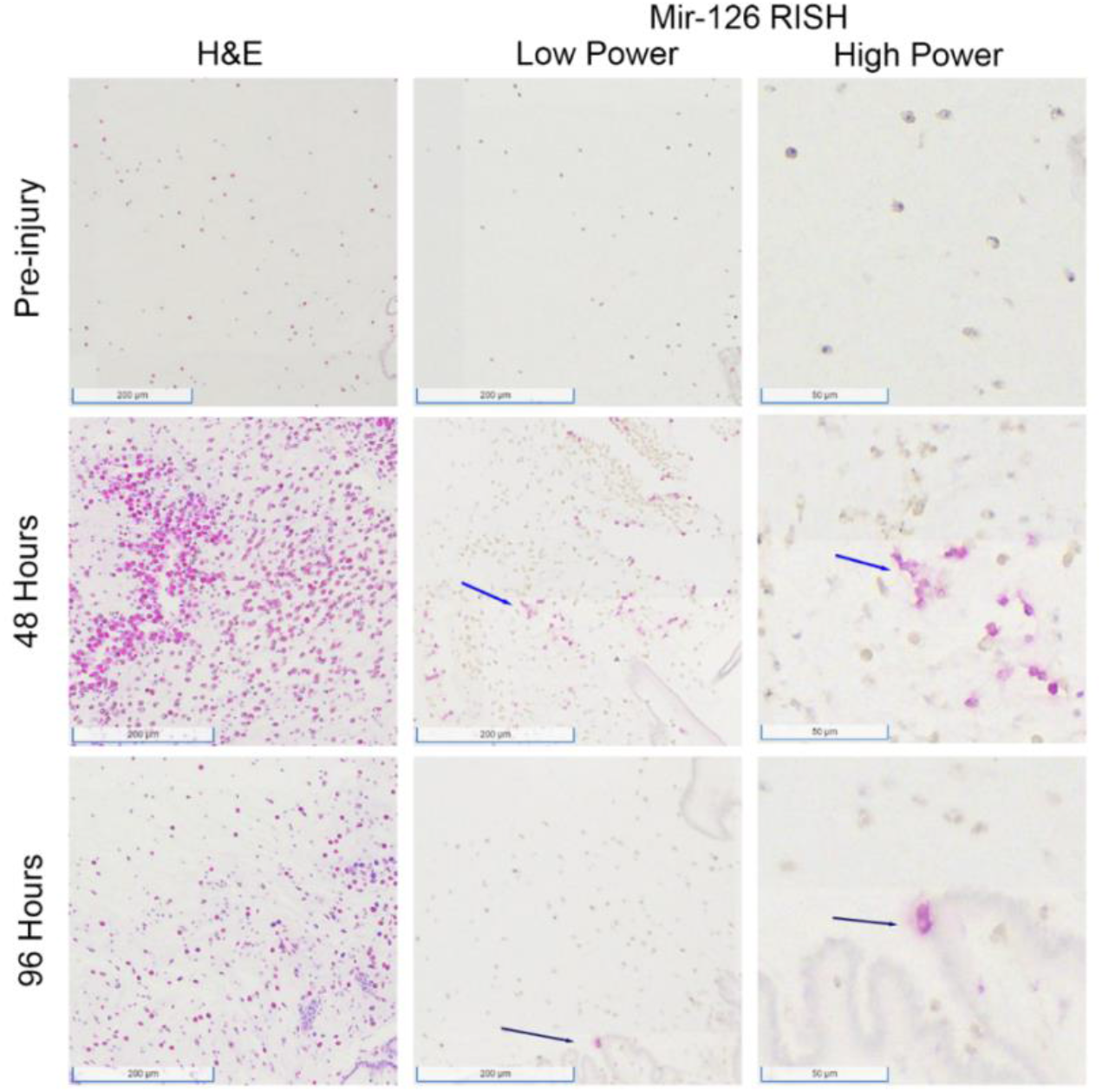
*C. Robusta* siphon injury model. Pre-injury there are few blood cells in the test near the siphon with none of these cells being Mir-126 positive. At 48 post amputation, there is a heavy infiltrate of blood cells in the test, a small subset of which are Mir-126 positive granular amebocytes. At 96 hours, the cellular infiltrate in the test is reduced and Mir-126 positive cells are rare.

MIR-126 and the endothelium provides a fascinating example of potential cell-type lineage phylogeny as this is the only known instance of where the evolution of the microRNA *precedes* the acquisition of the cell type. Mir-126 is specifically expressed in endothelial cells in Vertebrates, but this microRNA is also present in tunicates that lack endothelial cells. Taking advantage of the high conservation at the nucleotide level of the mature (3p) strand, we were able to employ RNA-in-situ-hybridization (RISH) from which we discovered that Mir-126 in *Ciona* is exclusively expressed in granular amebocytes, an invertebrate hemoblast cell. Our injury model did not induce upregulation of Mir-126 in other cell types. These findings support the long-proposed scenario that endothelial cells arose from hemoblasts and are related to thrombocytes, megakaryocytes, and their derivative platelets that characterize all extant vertebrates including the jawless fishes (Figure 6) (Casley-Smith, 1980; Fänge, 1998; Grant et al., 2017; Kuprijanov, 1990; Kuter et al., 1997; Munoz-Chapuli et al., 2005; Sabin, 1920). It further supports why Mir-126 is also found at lower levels in thrombocytes. Although complex transcriptional profiles of protein coding genes are known and can be used to distinguish between cells types (Wagner, 2014), in this case Mir-126 is far more informative given that the relevant protein-coding genes either evolved long before or coincident with the cell-type in question.

**Figure 6.**
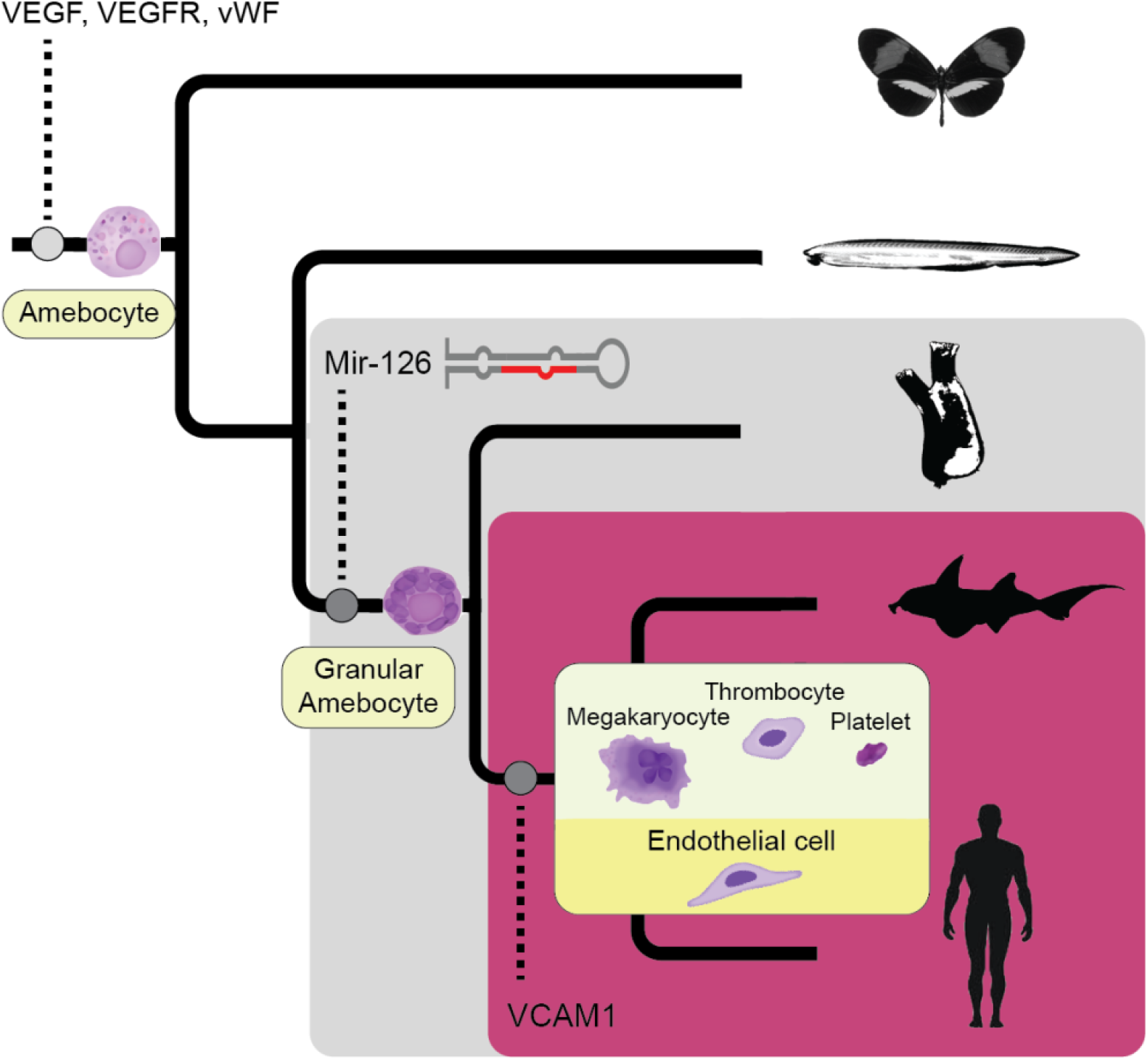
Cellular evolution of endothelial cells and thrombocyte. A primordial amebocyte is present before the rise of Metazoa, which contained VEGF, VEGFR and vWF. Through evolution, a granular amebocyte appeared. This is the first cell type to contain Mir-126 in tunicates. This cell evolved into endothelial cells, which, in this lineage, exclusively express VCAM1 and evolved into the related thrombocytes, megakaryocytes and platelets.

Like with most cell types, the initial specification of endothelial cells is not dependent on microRNA activity as Mir-126 knock-outs do not result in 100% embryonic lethality (Wang et al., 2008). Instead, a subset of Mir-126^-/-^ mice survive to adulthood, emphasizing the importance of robustness of microRNA activity on cell-type specification. Nonetheless, when the specification process does go awry in the absence of Mir-126 activity, the phenotypic consequences are profound with most mice dying from severe anemia (Fish et al., 2008; Kuhnert et al., 2008; Wang et al., 2008). Although initial knock-out studies incorrectly attributed this vascular defect to the host protein-encoding gene *Egfl7* (Schmidt et al., 2007), careful dissection of this two-gene locus whereby the knock-out of *Egfl7* without the perturbation of Mir-126 results in a phenotypically normal mouse (Fish et al., 2008; Kuhnert et al., 2008; Wang et al., 2008). Further, the introduction of a frame-shift mutation that results in the non-sense mediated decay of the egfl7 transcript in the zebrafish again results in a phenotypically normal fish (Rossi et al. 2015). These data suggest that the primary role of *Egfl7* is to simply serve as the transcriptional host for Mir-126. Indeed, the inhibition of *egfl7* transcriptional elongation results in a severe vascular phenotype similar to the Mir-126 knock-out phenotype (Rossi et al., 2015). It is likely then that the loss of the *Egfl* locus is manageable if the transcription of this now long non-coding RNA is not perturbed, possibly explaining the loss of *Egfl* loci across Olfactores evolution including not only tunicates, cyclostomes, but at least one population of *Homo sapiens* (Sulem et al., 2015).

This project provides new understanding of the evolution of Mir-126 in Olfactores and its key developmental role in the origin of new cell-types in the advent of vertebrates. The observed change of expression of Mir-126 from proto-endothelial granular amebocytes in the tunicate to endothelial cells in vertebrates is the first direct observation of the evolution of a cell-type in relation to microRNA expression indicating that microRNAs may indeed be a prerequisite of cell type evolution and can be used to trace the evolution of cell types. Despite this observation, there are notable limitations of this work. We have no clear idea of the regulatory roles of Mir-126 in *C. robusta* granular amebocytes. Careful studies in vertebrates have shown that there are two clear and phylogenetically conserved Mir-126_3p targets, the angiogenic repressors coded by Spred1 and Pik3r2 (Fish et al., 2008; Kuhnert et al., 2008; Wang et al., 2008). Each of these genes are one of three paralogues resulting from the 2R events acting on single gene loci (see Figure 2). Both vertebrate-specific Spred1 paralogues (Spred2 & 3, (Wang et al., 2008)) lack Mir-126 response elements, suggesting that the integration of Spred1 into the endothelial-specific gene regulatory network happened after the 1R event. Indeed, in *C. intestinalis* neither the *Spred* nor the *Pik3r* loci have a clear 7- or 8-mer Mir-126_3p binding site between the translational stop codon and the transcriptional start site of the next downstream (i.e., 3’) gene. Furthermore, current single cell RNA-seq data in *Ciona* are only available during developmental stages before these cells exist (Cao et al., 2019; Chacha et al., 2022). Therefore, we do not know what genes are expressed in granular amebocytes, and future functional studies of Mir-126 in *Ciona* will be needed to address this role.

In conclusion, we localized Mir-126, a microRNA expressed highly in endothelial cells and lowly in thrombocytes/platelets, to granular amebocytes in *C. robusta*. This, along with genomic evolution data, provides strong evidence for the speciation of endothelial cells and thrombocytes from a shared common ancestor cell and highlight the important role of microRNAs in cell evolution, furthering our understanding of evolutionary biology.

## ACKNOWLEDGEMENTS

*D. rerio* generously provided by Rebecca Rose of the Andrew McCallion lab. Digital cell images were created by Carlie Hruban and Josephine Halushka. We thank Norman Barker for photography assistance and Arun Patil for graphing assistance. This work was supported by the National Institute of General Medical Sciences, R01GM130564 and the National Heart, Lung, and Blood Institute, R01HL137811 to M Halushka.

## CONFLICT OF INTEREST

MKH is a consultant for Kiniksa Pharmaceuticals.

## DATA AVAILABILITY STATEMENT

There are no new datasets from this work.

**Supplementary Figure 1:** Gross dissection of *C. robusta*. **A**. A complete adult tunicate after being anesthetized (each square = 0.5”). **B**. A small tunicate processed whole. **C**. A vertical section of an adult tunicate. **D**. A sagittal section of an adult tunicate. **E**. Siphon area collected 48 hours post injury.

**Supplementary Figure 2:** Additional RISH of Mir-126, negative controls, H&E, and PAS staining of *C. robusta* **A**. H&E of the lymph organ (as seen in Fig 3). **B**. Mir-126 RISH of the lymph organ (as seen in Fig 3). **C**. Mir-126 negative control of the lymph organ. **D**. H&E of the test and cuticle. **E**. False positive Mir-126 RISH blush of the test and cuticle **F**. Negative control of test and cuticle. **G**. Mir-126 staining of mucous around an ovary. *H*. negative control of the ovary. **I**. H&E of the neural gland. **J**. Mir-126 RISH of the neural gland highlighting a serpentine pattern. **K**. Negative control. **L**. PAS stain highlighting the same serpentine pattern.

## Notes

### Competing Interest Statement

The authors have declared no competing interest.

