## Supplementary figures and images for "Direct observation of the evolution of cell-type specific microRNA expression signatures supports the hematopoietic origin model of endothelial cells"

### Supp_Fig_S1.png

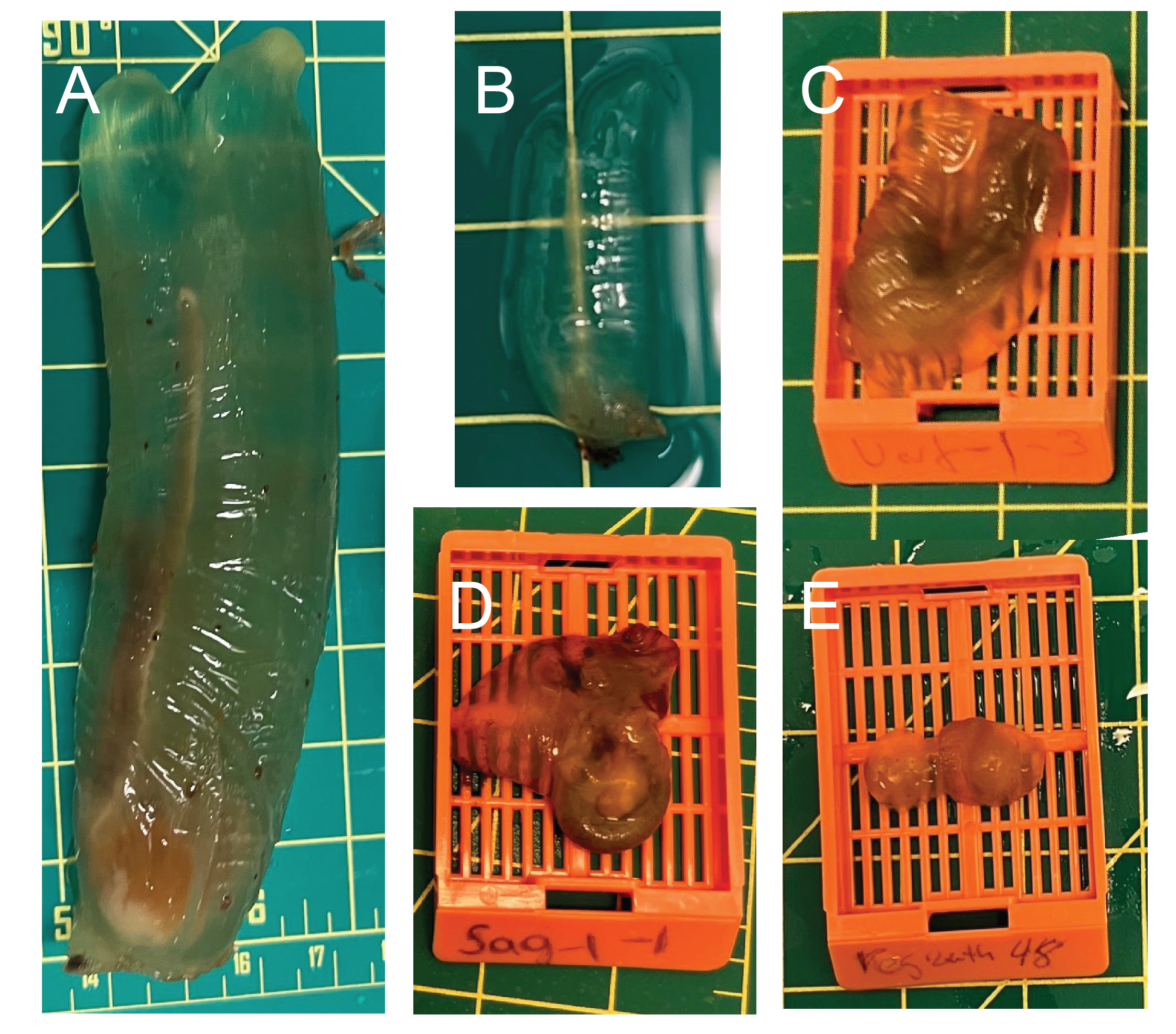

### Supp_Fig_S2.png

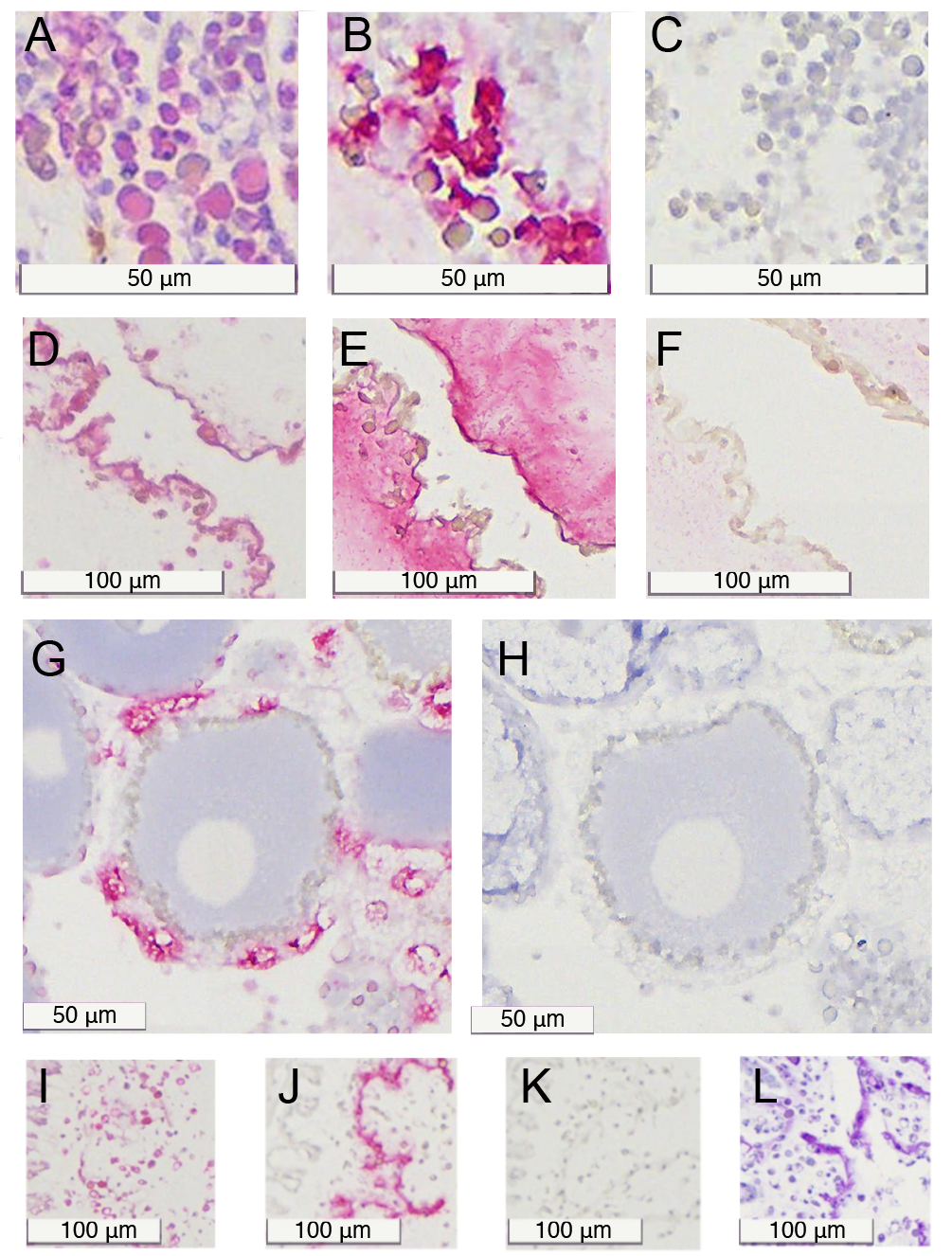
